# Cell competition for cancer treatment with Urine Progenitor Cells

**DOI:** 10.1101/2023.08.27.554858

**Authors:** Juli R. Bagó, Dhwani Radhakrishnan, Matouš Hrdinka, Tomasz Cichoń, Ryszard Smolarczyk, Sandra Charvátová, Benjamin Motais, Sebastian Giebel, Roman Hájek

## Abstract

Here, we provide evidence that urine progenitor cells (UPCs) might offer a novel therapeutic strategy for treating different types of cancer. We found that UPCs have inherent antitumor properties due to cell-cell competition, and we described their mechanism of action against different tumor cell lines. *In vitro* time-lapse analysis showed that the UPCs have tumor tropic properties, homing to various tumor-conditioned mediums. Besides, UPCs engineered for the expression of cytotoxic agents (the Tumor Necrosis Factor ligand superfamily member 10 and Herpesvirus-thymidine kinase) significantly increased their antitumor properties against several tumor cell lines and proved to serve as highly effective drug-delivery vehicles. Finally, using a mouse model of human triple-negative breast cancer, we observed a 150-fold decrease in tumor volumes in mice treated with engineered UPCs compared to controls. Collectively, our findings demonstrate for the first time the antitumor properties of the UPCs and form a foundation to continue exploring the new concept of cell-cell competition in progenitor cells to treat different types of cancer.

## 1. Introduction

Cancer cell-based therapy is a rapidly growing research field that utilizes allogeneic or autologous cells to fight cancer and relieve medical conditions. So far, the sources of therapeutic cells employed have been variable, ranging from immune cells to endothelial progenitor cells. Recent groundbreaking outcomes in treating distinct hematological malignancies with T-lymphocytes modified with chimeric antigen receptors (CARs) placed cancer cell-based therapies in the spotlight as a new pillar in cancer treatment. Still, for the time being, it is a fledgling therapy, and thus, there are several roadblocks to overcome, such as tumor escape, cell homing, effector toxicity, and universal access. Therefore, new therapeutic cell sources are sought out to circumvent these drawbacks.

Healthy and malignant tissues are highly functional networks of heterogeneous cell types with many complex interactions between neighboring cells. Cell competition is one of these intracellular interactions^[1-2]^, which participates in maintaining tissue and organ homeostasis. In this concept, the cells with higher fitness, named “winners,” eliminate surrounding cells with lower fitness, the “losers.” This phenomenon was first described in 1998 in Drosophila flies carrying the mutations in the ribosomal protein genes^[3]^. Later, similar processes of cell elimination by competition were observed in Drosophila wing discs overexpressing c-Myc^[4-5]^ and in mouse embryonic stem cells^[6-7]^. Interestingly, cells expressing higher levels of c-Myc (winners) in these models eliminated cells with lower c-Myc levels (losers). Thus, the expression levels of c-Myc could predict which neighboring cells would be removed through the development.

In various cell competition models, the clearance of less-fit cells can be achieved by different mechanisms. The most common mechanism is the induction of apoptosis in loser cells^[5, 8-10]^. In the epithelium, on the other hand, less-fit cells might be preferably eliminated by extrusion. Here, cells with oncogenic mutations, such as Trp53 and RasV12, are actively removed by wild-type epithelium cells^[11-14]^. This process of oncogenic cell extrusion, also known as the epithelial defense against cancer (EDAC), results in tumor cells’ death by necroptosis or anoikis. Finally, engulfing less-fit cells by surrounding healthy cells represents another described mechanism of cell elimination during cell competition^[6, 15]^.

We hypothesized that the phenomenon of cell fitness in cell-cell competition could be exploited for cancer treatment by selecting more-fit therapeutic cells to target less-fit tumor cells. To this end, due to their unique characteristics, we explored the so-called urine progenitor cells (UPCs), also known as urine stem cells or urine-derived stem cells. The UPCs are a subpopulation of urine-derived cells with stem cell properties, such as self-renewal and multipotential differentiation. Therefore, they express diverse gene markers that predict a strong fitness, such as the ones related to pluripotent stem cells^[17-18]^ (POU Class 5 Homeobox 1 (POU5F1), transcription factor 3 / 4 (Oct 3 / 4), stage-specific embryonic antigen 1/4 (SSEA-1/4), c-Myc and Kruppel-like factor 4 (Klf-4)), and those related to adult stem cells, such as CD90 and CD44. The UPCs can be isolated from regular voided urine via a non-invasive, simple, and low-cost approach and easily cultured *in vitro*, rapidly generating clinically relevant cell numbers in a short time^[16]^. Notably, another essential feature that makes UPCs an ideal cell source for therapeutic approaches is their safety, as there is no evidence of teratoma formation after *in vivo* transfusion of UPCs^[18]^.

## 2. Materials and Methods

### 2.1. UPCs Isolation, Expansion, and Cell Lines Cultivation

The UPCs were obtained from three voided urine sample donors (age range 34-46 from two males and one female). Voided urine samples (∼200 ml) were centrifuged at 300xg for 5 min. After discarding the supernatant, the cell pellet was resuspended and cultivated in a 24-well plate with a 1:1 medium of defined Keratinocyte-SFM (Thermo Fisher) and complete embryonic fibroblast medium (Cell Biologics). After 24 hours, the medium was changed to remove the unattached cells and detritus. Confluence was reached in 2-3 weeks. Every other day, the medium was replaced entirely. A limiting-dilution method was performed to study the proliferation capacity of a single UPC. To this end, 0.1 ml of a cell suspension with five UPC/ml was transferred in a 96-well plate to ensure a cell density as low as 0.5 cells per well. Seeding such an average of cells, and according to a Poison distribution, ensures that some wells will receive a single cell, reducing the probability of receiving more than one cell per well. Proliferation rates of the single-cell clones were monitored over time, and duplication time was calculated as follows:

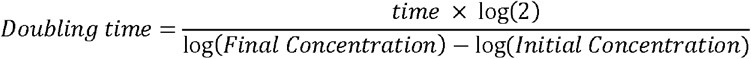

The panel of cancer cells consisted of the human lung carcinoma cell line A549 (a gift from Dr. Peter Dráber, BIOCEV, Prague, Czech Republic), the human osteosarcoma cell line U-2 OS (provided by Prof. Václav Hořejší, Institute of Molecular Genetics, Prague, Czech Republic), the human triple-negative breast cancer cell line MDA-MB-231 (ATCC HTB-26), the human ovarian adenocarcinoma OVCAR-3, the human prostate cancer cell line PC3 and the human colon cancer cell line HCT116 (all three cell lines provided by Dr. Ryszard Smolarczyk, Maria Skłodowska-Curie National Research Institute of Oncology, Gliwice, Poland).

All tumor cell lines, except for U-2 OS, were cultured in the RPMI 1640 medium (Sigma-Aldrich, Munich, Germany) supplemented with 10% of FBS (Sigma-Aldrich, Munich, Germany), 1% of penicillin-streptomycin and 1% of Ultraglutamine-1 (Lonza, Basel, Switzerland). U-2 OS cells were grown in high-glucose DMEM (Sigma-Aldrich, Munich, Germany) with 10% FBS and 1% penicillin-streptomycin (Lonza, Basel, Switzerland).

Human MSC derived from the cord blood (Promocell, Heidelberg, Germany) and human fibroblast cell line BJ-5ta (a gift from Dr. Raul Peña, Institut Hospital Mar Investigacions Mediques - Fundació IMIM, Barcelona, Spain) were cultured in high-glucose DMEM (Sigma-Aldrich, Munich, Germany) with 10% of FBS (Sigma-Aldrich, Munich, Germany) and 1% of penicillin-streptomycin (Lonza, Basel, Switzerland).

### 2.2. Phenotypic Analysis of UPCs

To evaluate phenotype markers’ expression on isolated and expanded UPC, we performed a flow cytometry analysis (BD FACSAria™ III Cell Sorter (BD Biosciences-US) with the following antibodies: CD44-FITC (Miltenyi, Bergisch Gladbach, Germany), CD90-APC (Miltenyi, Bergisch Gladbach, Germany), CD45-PB (Agilent Dako, Santa Clara, CA, USA) and CD34-APC (Miltenyi, Bergisch Gladbach, Germany).

### 2.3. Analysis of Antitumor Capacity of the UPCs

Antitumor assays were carried out in triplicates in flat white 96-well plates with a clear bottom (Thermo Scientific). Per well, 2000 tumor cells (target cells) tagged with luciferase expression were mixed with 2000 UPC (T:E = 1:1), 10000 UPC (T:E = 1:5) and 20000 (T:E = 1:10) in a total volume of 100 µl RPMI 1640 complete medium. 24 hours later, the total number of target cells was measured by adding D-luciferin potassium salt (Goldbio) to a final concentration of 0.5 mg/ml and reading the bioluminescence with the Infinite F Plex microplate reader (Tecan). The bioluminescence peak values were taken for each well, and relative cell viability was determined by dividing the registered photon with the corresponding photon counts from the control.

### 2.4. Migration Assay

The migration capabilities of UPCs were assessed in chemotaxis transwell assays with cancer cell lines serum-free conditioned media from MDA-MB-231, A549, and U-2 OS (48 hours). Transwell permeable supports (Costar) with 8 µm pores were placed in a 24-well plate with 0.5mL of conditioned media from the cancer cell lines. 0.2mL of medium without serum containing 0.5×10^5^ serum-starved (24 hours) UPCs were transferred to the upper chamber. 24 hours later, UPCs cells that migrated through the pores were collected from the bottom chamber and counted using the Luna-Stem Dual Fluorescence Cell Counter (Logos Biosystems).

### 2.5. Generation of Lentivirus Vectors, Production and Transduction

The lentiviral construct for the constitutive expression of the bioluminescence (firefly luciferase) and the red fluorescence (mCherry) was generated by amplifying the cDNA encoding both genes from Addgene Plasmid #44965 and cloned in the lentivirus backbone (Addgene #12262) using standard cloning procedures. The genes encoding TNFSF10 and TK were synthesized (GenScript, New Jersey, US) and cloned into the lentiviral backbone using standard cloning procedures. In the case of the TNFSF10 it was cloned with the extracellular domain of Flt3L to ensure it secretion.

The lentivirus constructs were packaged as a lentivirus vector in human embryonic kidney 293FT cells (HEK 293FT, a kind gift of Prof. Vaclav Hořejší, Institute of Molecular Genetics, Prague). Briefly, 293FT cells were seeded at 3 x10^6^ cells per 10 cm dish in a complete DMEM medium. The next day, the cells were transfected with the plasmid encoding the gene interest, the pMD2.G plasmid encoding the VSV-G envelope (Addgene #12259), and the psPAX2 packaging plasmid (Addgene #12260) using the jetPRIME transfection system (Polyplus). The medium was replaced four hours after transfection, and the viral supernatant was collected three days later. The viral supernatant was filtered with a 0.45 µm filter and concentrated with Amicon Ultra-15 filter tubes (Millipore). The tumor cell lines and UPCs were infected with the concentrated virus in a culture media containing 8 µg/ml Polybrene (Sigma) and positively transduced cells selected by FACS for the expression of the corresponding fluorescent reporter.

### 2.6. Generation of Knockout with CRISPR/Cas9

The CRISPR-Cas9 technology of two vector systems was applied to generate the knockouts. The plasmids for transfection were purchased from Addgene (Lenti-Cas9 Blast #52962 and Lenti-guide Puro, #52963). The guide RNAs were outlined using https://chopchop.cbu.uib.no/ website. Three guide RNAs (sgRNA) were purchased per gene and cloned into the Lenti-guide Puro vector using standard cloning procedures. The mixture of three lentivirus constructs was packaged as a lentivirus vector in human embryonic kidney 293FT cells as described above. The UPCs, MDA-MB-2331, and U-2 OS were transduced with the different lentivirus vectors in culture media containing 10 µg/mL of Polybrene (Sigma). After seventy-two hours of infection, transduced cells were selected by blasticidin (10µg/ml) and puromycin (1µg/ml) resistance. The selected cells were expanded and used for experiments. The efficiency of knockout generation was assessed by SYBR green qRT-PCR method.

### 2.7. In vivo Experiments

All animal-related procedures were carried out with the approval of the Local Ethics Commission of Animal Experiments in Katowice (permission No: 48/2021) at the Maria Sklodowska-Curie National Research Institute of Oncology in Gliwice, Poland. Mice were housed in a pathogen-free facility in SPF standard in a HEPA-filtered Allentown’s IVC System (Allentown Caging Equipment Co, NJ, USA). The animals received a total pathogen-free standard diet (Altromin 1314, Altromin Spezialfutter GmbH & Co. KG, Germany) and water ad libitum throughout the whole study. Animals were treated in accordance with the recommendations in the Guide for the Care and Use of Laboratory Animals of the National Institutes of Health. Experiments on animals were conducted in accordance with the 3R rule.

Adult 6–8-week-old female severe combined immunodeficient mice (SCID) were ordered from Charles River Laboratories (Wilmington, MA, USA). After 14 days of acclimatization, the mice were divided into two groups: the treated group that received the rear flank subcutaneous injection of 100 µL of PBS containing 2 × 10^6^ MDA-MB-231 and 2 × 10^6^ UPCs-TNFSF10 mixed with 100 µL of Matrigel (Corning, Corning, NY, USA) and the control group, with 4 SCID mice that received the rear flank subcutaneous injection of 100 µL of PBS containing 2 × 10^6^ MDA-MB-231 mixed with 100 µL of Matrigel (Corning, Corning, NY, USA). Tumor progression was measured with calipers, and tumor volumes were calculated using the following formula: volume = width^2^ × length × 0.52. Mice with considerable weight loss (>20% of their initial body weight) or, with signs of pain (rough hair coat or hunched posture), or with tumors exceeding 2000 mm^3^ were sacrificed. On day 25 after implantation, mice from the different groups were sacrificed by cervical dislocation, and tumors were retrieved.

## 3. Results and Discussion

First, to evaluate the feasibility of generating clinically relevant cell numbers, we isolated and expanded UPCs from several healthy donors as described in the methods section, with an average of four UPCs from 200 ml voided urine. We studied the proliferative capacity of single UPC clones over time by applying the limiting-dilution method, obtaining a median duplication time of 27 hours in their exponential growth phase and a median of 2×10^5^ cells per clone (data not shown). Expanded UPCs were characterized by specific cell surface markers, such as the positive expression of stem cell markers CD44 and CD90 and the negative expression of hematopoietic stem cell markers CD34 and CD45 (**Fig. 1a**) ^[16, 19]^.

**Figure 1.**
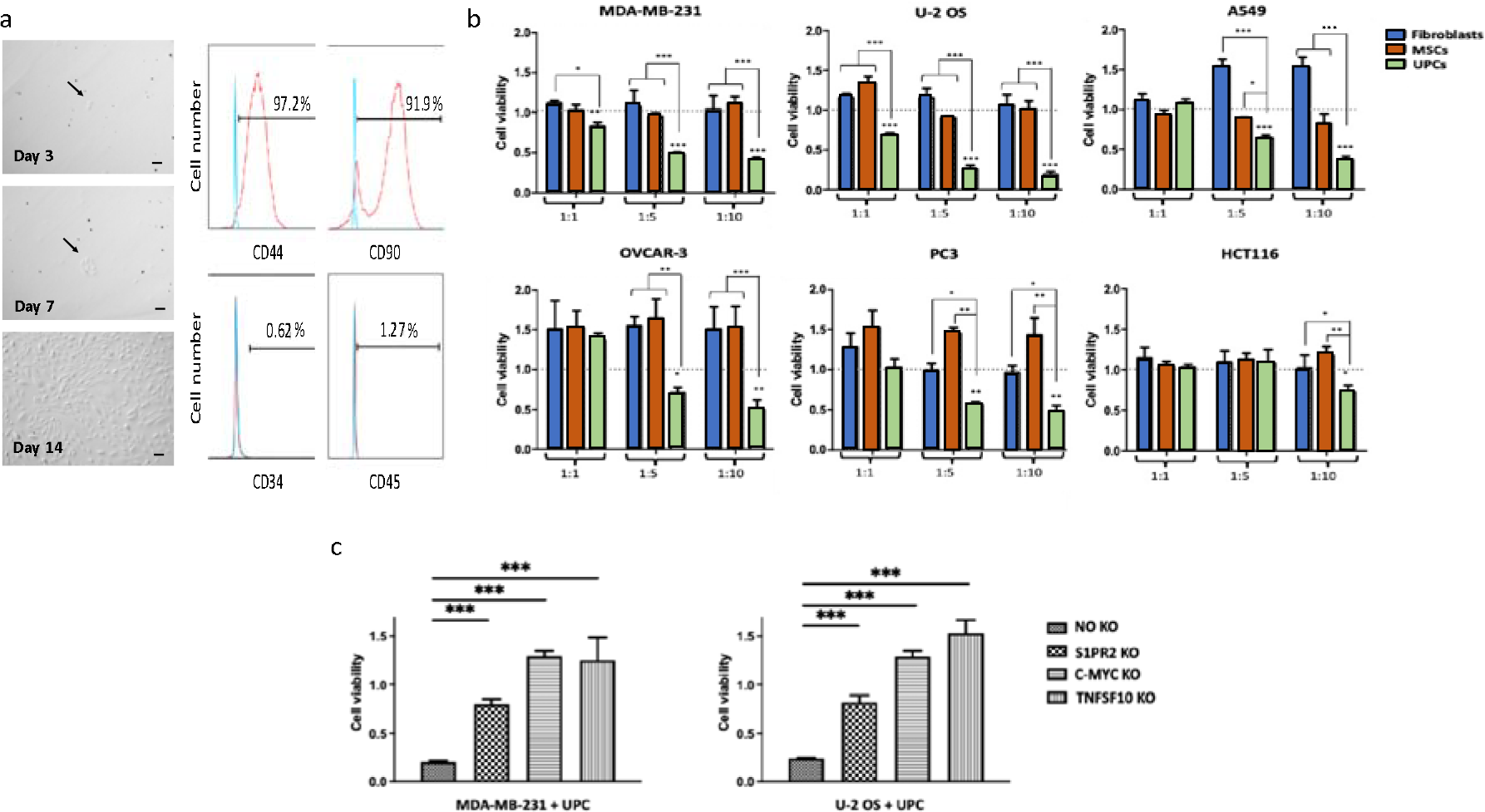
Characterization and antitumor properties of the UPCs. **(a) UPCs under an inverted microscope at different time points after isolation. Flow cytometry analysis of UPCs showing the expression of** CD44, CD90, CD34, and CD45 markers. (b) Summary graph showing cell viability of different tumor cell lines (U-2 OS, MDA-MB-231, A549, OVCAR-3, PC3, and HCT116) after coculture with Fibroblasts, MSC, and UPCs at different target-effector (T:E) ratios for 72 hours. Results are presented as a relative cell viability compared to the control (tumor cells without fibroblasts, MSC, and UPCs). (c) Viability of tumor cell lines MDA-MB-231 and U-2 OS after coculture with knockout UPCs for the genes S1PR2, C-MYC, and TNFSF10, at a T:E ratio of 1:10 for 72 hours. No KO refers to the cell line transduced with the control vector. Results are presented as relative cell viability compared to the control (tumor cells without UPCs). All results are representative of several (N=3) independent samples. Data are presented as means ± SD from three technical replicates. *, p < 0.05, **, p < 0.01, ***, p < 0.001, one-way ANOVA, repeated measures test. Scale bar 50 µm.

Next, we investigated the UPCs’ ability to mediate antitumor activity in cocultivation assays with a broad spectrum of tumor cells (U-2 OS, MDA-MB-231, A549, OVCAR-3, PC3, and HCT116) at different target (tumor cell)-effector (UPCs) (T:E) ratios for 72 hours. In all cases, we observed significantly lower numbers of tumor cells in cocultures with UPCs compared to control cultures without UPCs. Moreover, the tumor cell numbers in cocultures progressively decreased with increasing T:E ratios (**Fig. 1b)**.

To further evaluate the distinctive antitumor properties of the UPCs, we performed the same cocultivation assays with mesenchymal stem cells (MSCs) derived from the cord blood and fibroblast cell line BJ-5ta (fibroblasts) instead of UPCs. Interestingly, in these experiments, no significant reduction in tumor cell numbers was registered at any T:E ratio studied. On the contrary, a significant increase in the tumor cell numbers was observed in the cocultures of A549 with fibroblast, PC-3 with MSCs, and OVCAR-3 with fibroblasts and MSCs (**Fig. 1b**). Additionally, to test the antitumor specificity of the UPCs, the UPCs were cultivated with fibroblast at different ratios for 72 hours. In this case, the viability of fibroblasts was not significantly reduced at any of the T:E ratios tested compared to fibroblast cultures without UPCs **(Supplementary Fig. 1)**.

**Figure S1.**
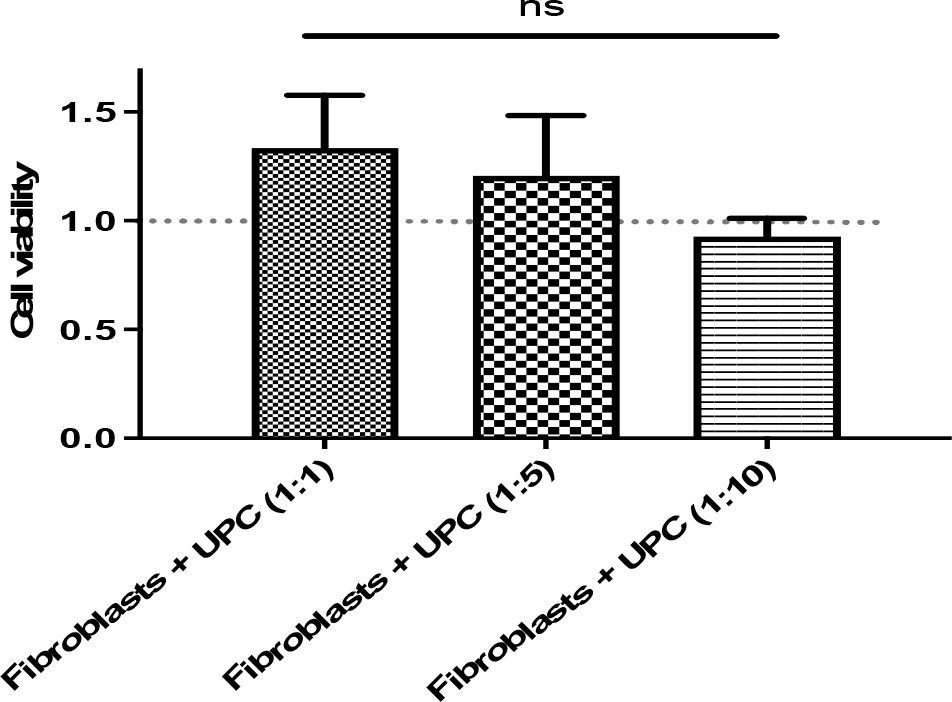
Summary graph showing the cell viability of fibroblasts after coculture with UPCs at different ratios for 72 hours. Results are shown as relative cell viability compared to the control (fibroblasts without UPCs). Results are representative of several (N=3) independent samples. Data are presented as means ± SD from three technical replicates. ns, not significant, one-way ANOVA, repeated measures test.

To the best of our knowledge, these results demonstrate the unique UPCs’ antitumor capacity towards different tumor cell lines for the first time.

The capacity of the UPCs to migrate towards the tumor target cells is paramount for an effective antitumor response. For this reason, we evaluated UPCs’ migration capacity using a Transwell migration assay, in which the UPCs were cultivated in an upper chamber, and their migration across a permeable support membrane toward the lower chamber with tumor cell-conditioned media was monitored over time. We observed a significant increase in the UPCs’ migration towards all the tumor cell-conditioned media tested (**Supplementary Fig. 2**), demonstrating their tumor tropic properties.

**Figure S2.**
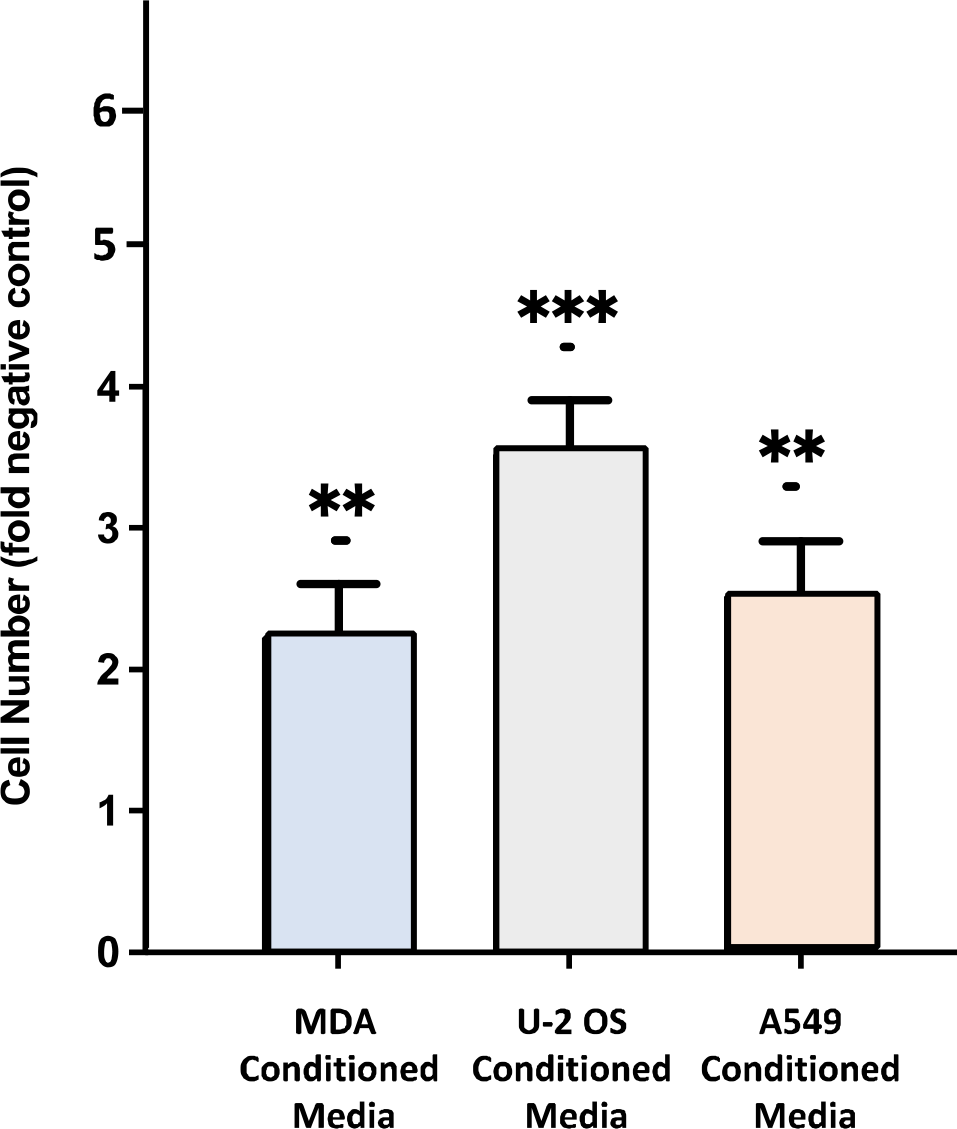
Transwell migration assay of UPCs to evaluate their ability to move towards conditioned media from U-2 OS, MDA MB-231, and A549. Bar graphs show a significant increase in UPCs migration towards conditioned media of MDA-MB-231, U-2 OS, and A549 compared to a negative control without conditioned media. Results are representative of several (N=3) independent samples. Data are presented as means ± SD of three technical replicates. *, p < 0.05, **, p < 0.01, ***, p < 0.001, Student’s t-test.

Since the UPCs express diverse markers that predict strong cell fitness, we hypothesized that the observed antitumor mechanisms of the UPCs might be linked to cell competition. To test our hypothesis, we conducted a series of *in vitro* experiments. First, we performed an experiment to reveal whether the UPC’s antitumor effect depended on cell-to-cell contact. To this end, the UPCs and tumor cells were cultivated in the same well but physically separated by a small pore-size membrane that only permits the diffusion of soluble factors, thus preventing direct contact between both cell populations. However, in this experiment, no significant differences in tumor cell viability were registered, suggesting the need for direct cell-to-cell interactions to observe the antitumor effect of UPCs (**Supplementary Fig. 3)**.

**Figure S3.**
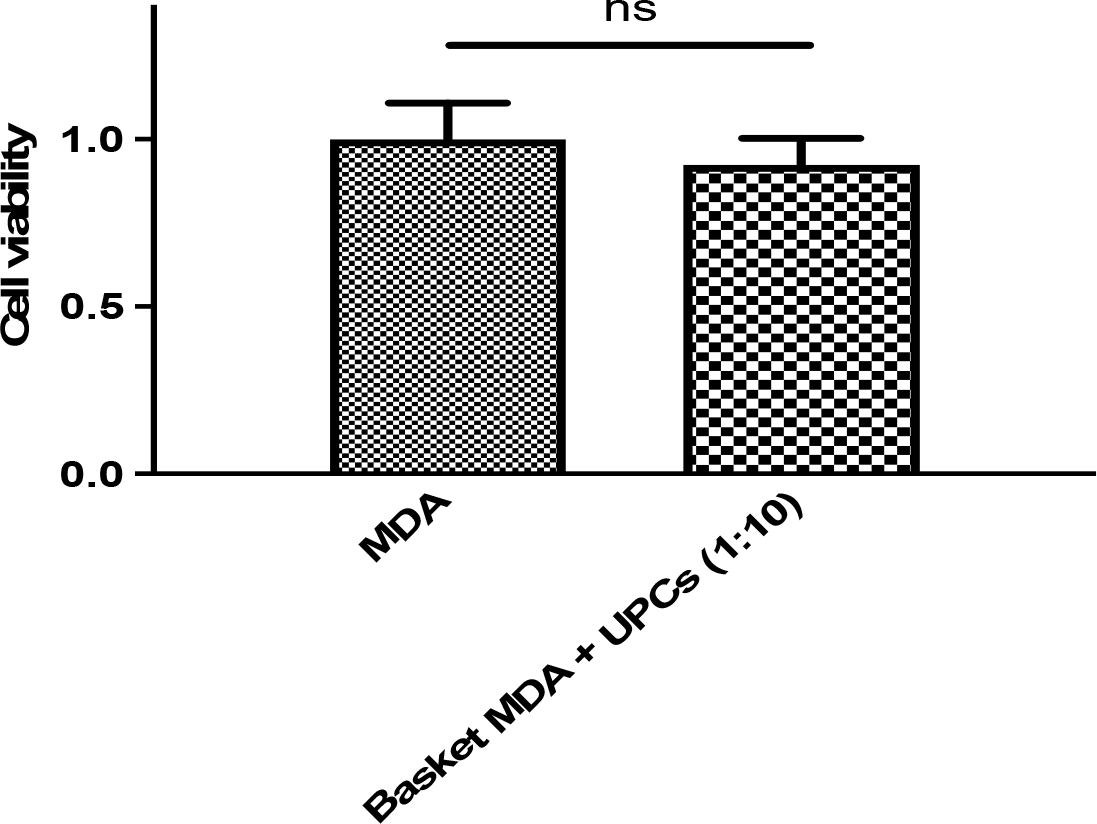
Cell viability of MDA-MB-231 cultivated with UPCs and separated by a transwell permeable support. Histogram with the MDA MB-231 viability after cultivation with UPCs separated by a small-sized pore membrane for 72 hours at ratio of 1:10 (T:E), compared to control without UPCs. Data are represented as a relative cell viability compared to the control (MDA without UPCs). No significant reduction in tumor cell viability was observed in these conditions. Results are representative of several (N=3) independent samples. Data are presented as means ± SD of three technical replicates. ns, not significant, Student’s t-test.

Once we ascertained that the UPC’s antitumor response depends on direct cell-to-cell contacts, we wanted to identify genes expressed in UPCs capable of triggering cell competition responses towards three main known mechanisms: apoptosis, phagocytosis, and extrusion. To this end, we evaluated the expression of genes involved in cell competition mechanisms using publicly available differential gene expression datasets, where the gene expression profiles of the UPCs were compared to MSCs and fibroblasts^[20-22]^. Interestingly, we found that UPCs overexpress three genes related to the antitumor response through cell competition. First, the Tumor Necrosis Factor ligand superfamily member 10 (TNFSF10/TRAIL), which binds to TNFRSF10A/TRAILR1, TNFRSF10B/TRAILR2, TNFRSF10C/TRAILR3, and TNFRSF10D/TRAILR4 in tumor cells and triggers their apoptosis^[23-24]^. Second, the transcription factor c-Myc (C-MYC) is a critical protein orchestrating cell competition and a reliable fitness indicator^[6-7]^. Lastly, the Sphingosine-1-Phosphate Receptor 2 (S1PR2) participates in the epithelial defense against cancer by the apical extrusion of tumor cells^[27]^.

To determine the relative implication of these genes in the UPCs-mediated cell-cell competition, we evaluated the antitumor properties of UPCs after knocking out the corresponding genes via lentiviral CRISPR/Cas9 genome editing. Successful gene deletions (decrease in gene expression levels) were confirmed by qRT-PCR **(Supplementary Fig. 4)**.

**Figure S4.**
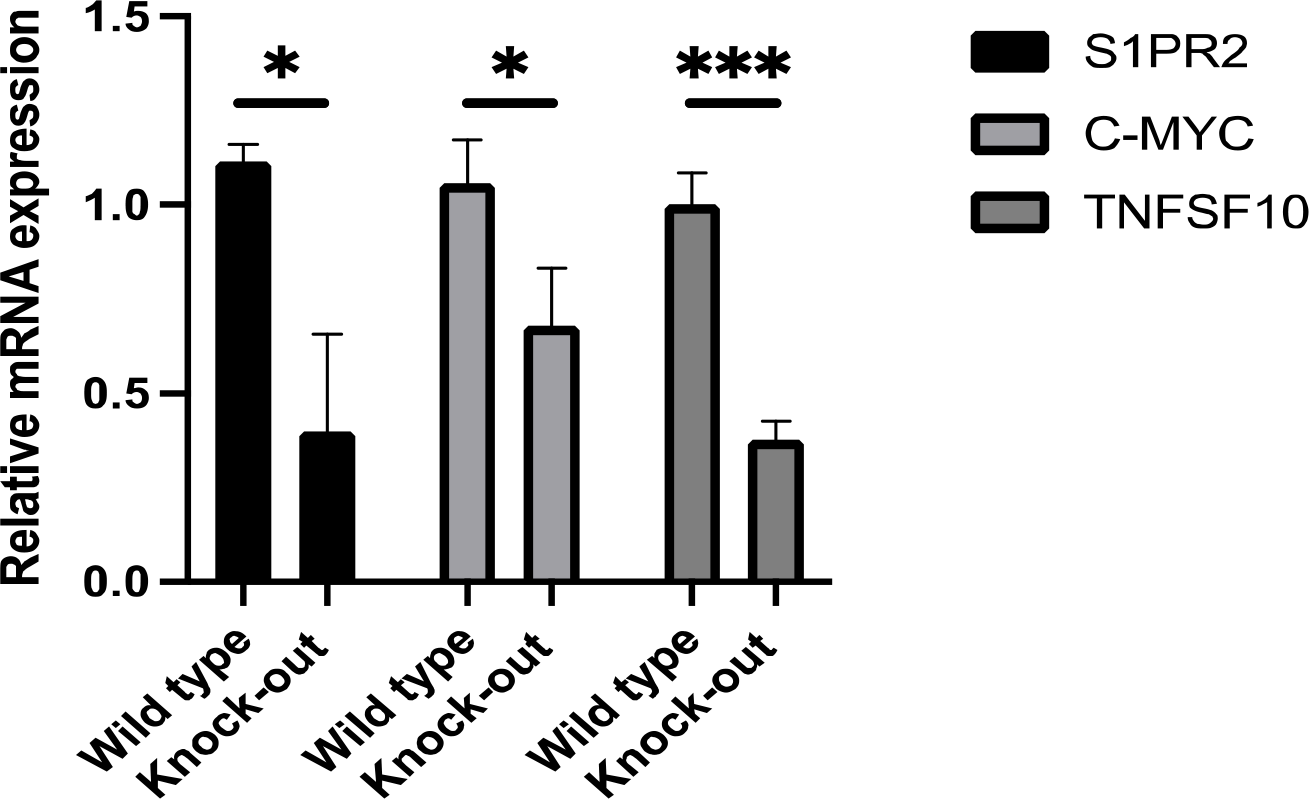
qRT-PCR analysis of relative expression levels of S1PR2, C-MYC, and TNFSF10 mRNA in wild-type cells compared to corresponding knockouts. Results are representative of several (N=3) independent samples. Data are presented as means ± SD of three technical replicates. *, p < 0.05, **, p < 0.01, ***, p < 0.001, Student’s t-test.

Decreased levels of C-MYC and TNFSF10 expression resulted in a complete abrogation of the antitumor properties of the UPCs towards the tumor cell lines U-2 OS and MDA-MB-231. On the other hand, decreased expression levels of the extrusion mediating gene S1PR2 in UPCs reduced the UPCs’ antitumor effect against tumor cell lines U-2 OS and MDA-MB-231 to more than 50% (**Fig. 1c)**. Collectively, our results indicate that the antitumor effects of UPCs depends on cell-cell contacts and are likely induced by cell competition, with three significant genes involved: C-MYC, TNFSF10, and S1PR2.

Next, we wanted to understand the cell death mechanisms induced in tumor cells during the competition with UPCs. In apoptosis, proteases called caspases play pivotal roles in activating and mediating cell death through proteolytic cascades. Two main pathways of triggering apoptosis rely on the initiator Caspase-8 (CASP8, extrinsic pathway) and Caspase-9 (CASP9, intrinsic pathway). To this end, we generated both CASP8 and CASP9 knockouts by CRISPR/Cas9 genome editing in MDA-MB-231 and U-2 OS cell lines and measured their viability after coculture with UPCs. Successful gene deletions were confirmed by qRT-PCR **(Supplementary Fig. 5**).

**Figure S5.**
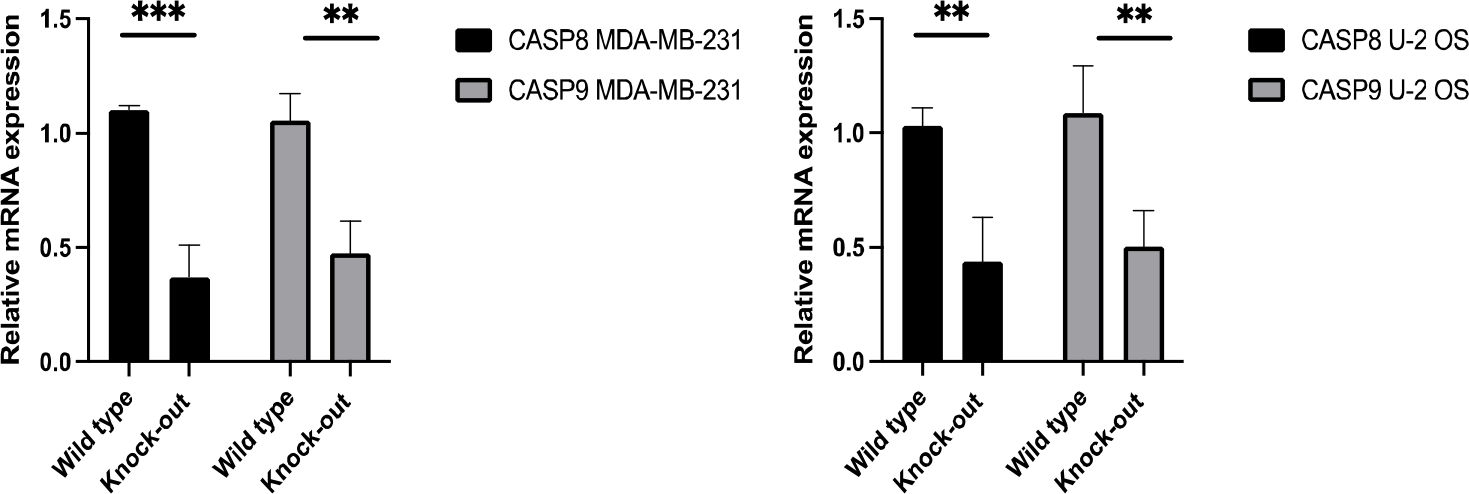
qRT-PCR analysis of relative expression levels of CASP8 and CASP8 mRNA in wild-type MDA-MB-231 cells and U-2 OS compared to corresponding knockouts. Results are representative of several (N=3) independent samples. Data are presented as means ± SD of three technical replicates. **, p < 0.01, ***, p < 0.001, Student’s t-test.

Interestingly, knocking out either CASP8 or CASP9 in MDA-MB-231 and U-2 OS cells significantly reduced the antitumor properties of the UPCs as compared to cocultures of UPCs with wildtype MDA-MB-231 and U-2 OS cells (**Fig. 2a)**.

**Figure 2.**
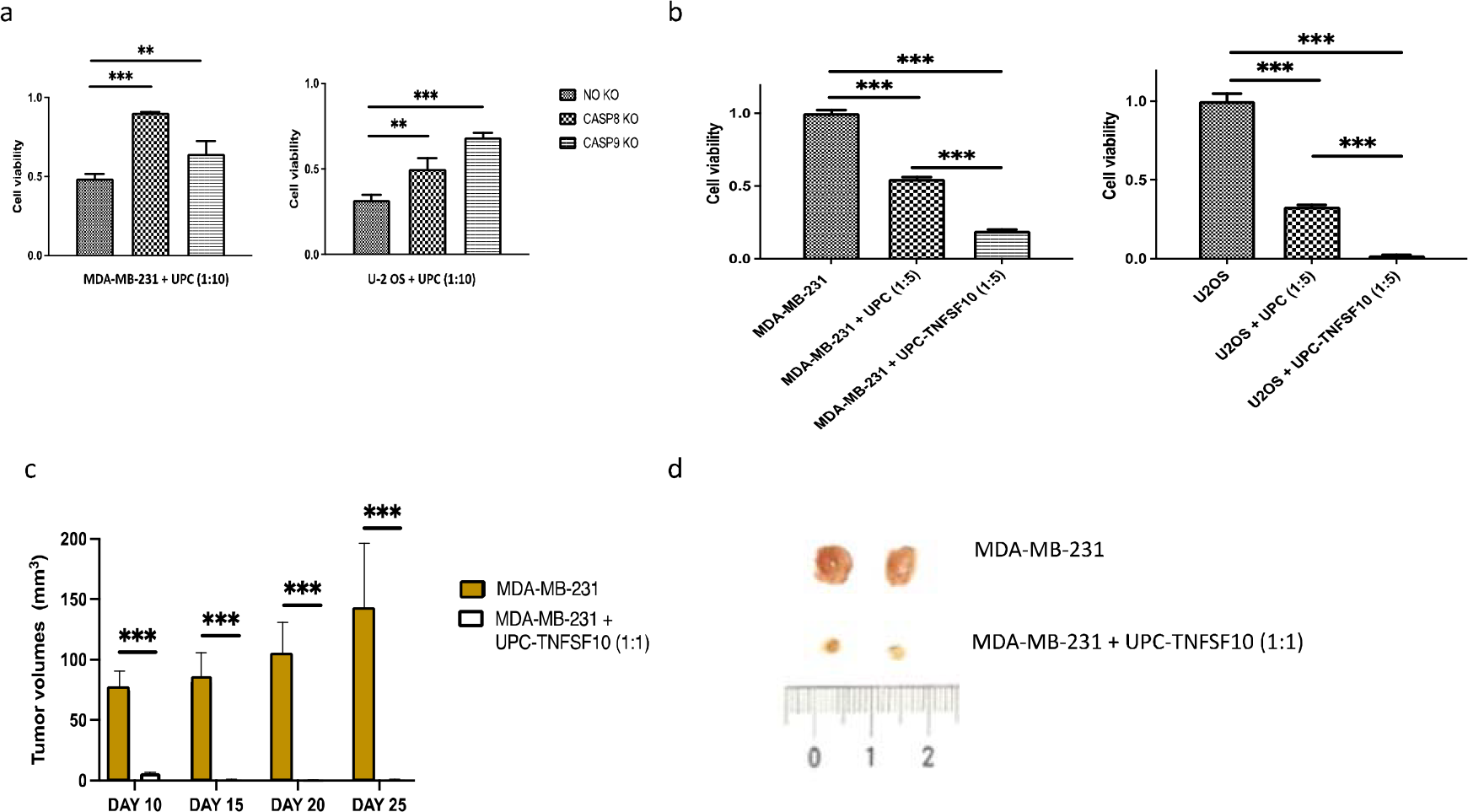
Apoptosis activation in tumor cell lines as a competition mechanism of UPCs and increase of UPCs antitumor properties after engineering for the overexpression of the apoptotic inducer TNFSF10, and its effectivity *in vivo*. **(a)** Graphic with tumor cell lines MDA-MB-231 and U-2 OS viability with knockout for the genes CASP8 and CASP9 after coculture with UPCs at a T:E of 1:10 for 72 hours. No KO refers to cell line transduced with the control vector. Data are represented as relative cell viability in comparison with the control (tumor cell lines without UPCs) (**b**) Summary graph showing the tumor cell viability of MDA-MB-231 and U-2 OS after coculture with UPCs engineered for the expression of TNFSF10 at a T:E of 1:5 for 72 hours, and compared to no engineered UPCs and control without UPCs. Data are represented as relative cell viability in comparison with the control (tumor cell lines without UPCs) (c) Summary graph of tumor volumes in the group of mice with MDA-MB-231 (*n* = 4) and a group of mice with MDA-MB-231 + UPC-TNFSF10 (*n* = 4) xenografts over time. (d) Representative images of tumors retrieved on day 25 after implantation from the two groups. Data in (a and b) represent of several (N=3) independent samples and are presented as means ± SD from three technical replicates. Data in (c) are presented as means ± SD from n = 4 for MDA-MB-231 and MDA-MB-231 + UPC-TNFSF10 groups. *, p < 0.05, **, p < 0.01, ***, p < 0.001, one-way ANOVA, repeated measures test, and Student’s *t*-test.

Collectively, these results strongly suggest that the induction of apoptosis signaling pathways is the primary mechanism used by UPCs to trigger cell death in tumor cells.

So far, we have shown that the UPCs display potent tumor cell-killing properties; however, the tumor cell elimination in coculture-killing assays was not complete even with the highest T:E ratios. Therefore, we attempted to increase the natural antitumor properties of the UPCs by gene engineering. In one approach, we engineered UPCs to over-express one of the main proteins involved in the UPCs cell competition against tumor cells, the TNFSF10/TRAIL. Because of its ability to selectively induce apoptosis in tumor cells, the TNFSF10/TRAIL protein is qualified as a potential drug specific for several different cancer types^[29]^. Indeed, the cocultivation of UPCs overexpressing TNFSF10 with tumor cell lines MDA-MB-231 and U-2 OS induced an even more robust anti-tumor response than non-engineered wild-type UPCs (**Fig. 2b**).

In the second approach to artificially boost the antitumor properties of UPCs, we evaluated the possibility of engineering UPCs for the constitutive expression of the Herpesvirus-thymidine kinase (HSV-TK) enzyme. Mechanistically, HSV-TK phosphorylates the nontoxic prodrug ganciclovir (GCV), converting it into a lethal drug that kills rapidly dividing neighboring cells by a bystander effect^[28]^. When we cultivated the tumor cell line U-2 OS with UPCs expressing HSV-TK (UPC-TK) in the presence of GCV, we noticed a significant reduction in tumor cell numbers compared to the cultivation of U-2 OS with non-engineered UPCs **(Supplementary Fig. 6**).

**Figure S6.**
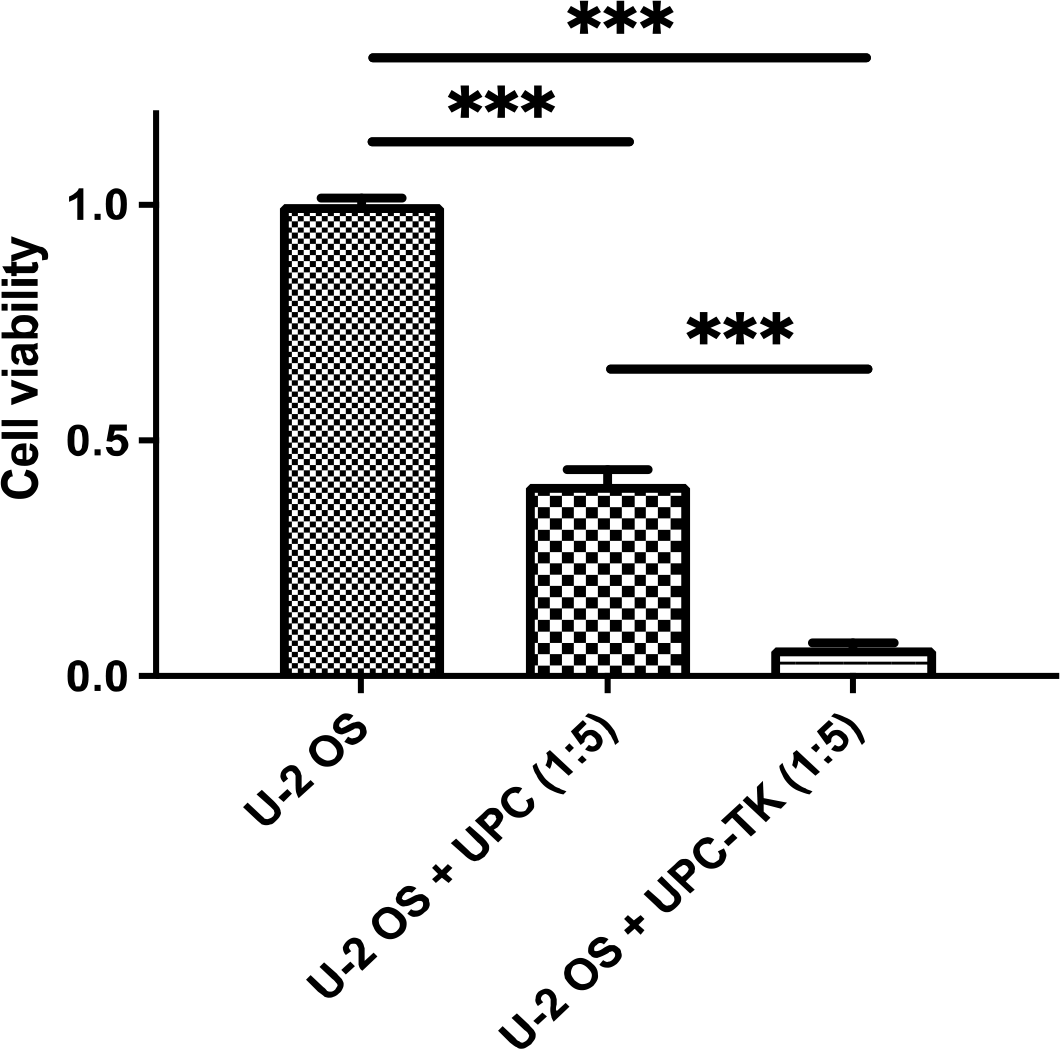
Antitumor properties of UPCs after engineering for the expression of HSV-TK. Graph with the tumor cell viability of U-2 OS after cultivation with UPCs-TK and GCV at a T:E ratio of 1:5 for seventy-two hours, compared to wild-type UPCs and control without UPCs. Results are presented as relative cell viability compared to the control (U-2 OS without UPCs). Results are representative of several (N=3) independent samples. Data are presented as means ± SD from three technical replicates. *, p < 0.05, **, p < 0.01, ***, p < 0.001, one-way ANOVA, repeated measures test.

Collectively, our results demonstrate the feasibility and efficiency of engineering the UPCs to increase their inherent antitumor properties with distinct cytotoxic agents.

Lastly, we investigated the *in vivo* efficacy of UPCs engineered to overexpress TNFSF10/TRAIL in a SCID mouse model of triple-negative human breast cancer. To this end, we injected human MDA-MB-231 subcutaneously in SCID mice with UPCs-TNFSF10/TRAIL or without UPCs as control and tracked tumor volumes over time. We observed that mice with UPCs-TNFSF10/TRAIL originated significantly smaller tumors than those without UPCs, with tumor volumes on day twenty-five after implantation 150-fold smaller than controls (**Fig. 2c-d**). Thus, our results suggest that the UPCs-TNFSF10/TRAIL are highly efficient in reducing the in vivo tumor growth of the breast cancer tumor cell line MDA-MB-231.

In summary, our results provide the first evidence that UPCs have inherent antitumor properties against different tumor types triggered by the cell-cell competition phenomenon. Besides, we found that the UPCs have tumor cell tropism, and they can be engineered with different cytotoxic agents to increase their inherent antitumor properties *in vitro* and *in vivo*. The presented data support the potential of UPCs to provide a viable alternative in cancer cell-based therapy, with their unique antitumor properties and capacity to serve as a highly effective drug-delivery vehicle. Together, these results form a foundation to continue exploring the new concept of cell-cell competition in progenitor cells to treat different types of cancer.

## Author Contributions

JRB., R.S., T.C., SC., and BM performed and analyzed the *in vitro* and *in vivo* data. D. R. and M.H., performed and analyzed the CRIPSR/Cas9 experiments. S.G., and R.H., assisted in the revision and editing of the manuscript. J.R.B. supervised and designed the whole work and wrote the manuscript.

## Institutional Review Board Statement

The study was conducted according to the guidelines of the Declaration of Helsinki, and approved by the Ethics Committee at the Faculty of Medicine in Ostrava (139/2023). All animal-related procedures were performed with the approval of the Animal Care Committee at the Maria Sklodowska-Curie National Research Institute of Oncology in Gliwice, Poland (48/2021).

## Acknowledgments

We thank Jan Vrána, Lucie Cerná and Veronika Kapustová for insightful work in cell analysis by FACS (Flow cytometry facility University Hospital of Ostrava).

## Data availability

All relevant data are within the manuscript and its Supporting Information files.

